# Sparse-Coding Variational Auto-Encoders

**DOI:** 10.1101/399246

**Authors:** Victor Geadah, Gabriel Barello, Daniel Greenidge, Adam S. Charles, Jonathan W. Pillow

**Author notes:** These authors contributed equally: Victor Geadah, Gabriel Barello.

## Abstract

The sparse coding model posits that the visual system has evolved to efficiently code natural stimuli using a sparse set of features from an overcomplete dictionary. The original sparse coding model suffered from two key limitations, however: (1) computing the neural response to an image patch required minimizing a nonlinear objective function via recurrent dynamics; (2) fitting relied on approximate inference methods that ignored uncertainty. Although subsequent work has developed several methods to overcome these obstacles, we propose a novel solution inspired by the variational auto-encoder (VAE) framework. We introduce the sparse-coding variational auto-encoder (SVAE), which augments the sparse coding model with a probabilistic recognition model parametrized by a deep neural network. This recognition model provides a neurally plausible feedforward implementation for the mapping from image patches to neural activities, and enables a principled method for fitting the sparse coding model to data via maximization of the evidence lower bound (ELBO). The SVAE differs from standard VAEs in three key respects: the latent representation is overcomplete (there are more latent dimensions than image pixels), the prior is sparse or heavy-tailed instead of Gaussian, and the decoder network is a linear projection instead of a deep network. We fit the SVAE to natural image data under different assumed prior distributions, and show that it obtains higher test performance than previous fitting methods. Finally, we examine the response properties of the recognition network and show that it captures important nonlinear properties of neurons in the early visual pathway.

## 1 Introduction

Generative models have played an important role in computational neuroscience by offering normative explanations of observed neural response properties (Olshausen & Field, 1996a,b, 1997; M. Lewicki & Olshausen, 1999; Dayan et al., 2003; Berkes & Wiskott, 2005; Coen-Cagli et al., 2012). These models seek to model the distribution of stimuli in the world *P*(**x**) in terms of a conditional probability distribution *P*(**x**|**z**), the probability of a stimulus **x** given a set of latent variables **z**, and a prior over the latent variables *P*(**z**). The advantage of this approach is that it mimics the causal structure of the world: an image falling on the retina is generated by “sources” in the world (e.g., the identity and pose of a face, and the light source illuminating it), which are typically latent or hidden from the observer. Perception is naturally formulated as the statistical inference problem of identifying the latent sources that generated a particular sensory stimulus (D. Knill & Richards, 1996; Weiss et al., 2002; D. C. Knill & Pouget, 2004; Moreno-Bote et al., 2011). Mathematically, this corresponds to applying Bayes’ rule to obtain the posterior over latent sources given sensory data: *P*(**z**|**x**) ∝ *P*(**x**|**z**)*P*(**z**), where the terms on the right-hand-side are the likelihood *P*(**x**|**z**) and prior *P*(**z**), which come from the generative model.

Perhaps the famous generative model in neuroscience is the *sparse coding model*, introduced by Olshausen and Field (Olshausen & Field, 1996a,b) to account for the response properties of neurons in visual cortex. The sparse coding model posits that neural activity represents an estimate of the latent features underlying a natural image patch under a linear generative model. The model’s key feature is sparsity: a heavy-tailed prior over the latent variables ensures that neurons are rarely active, so each image patch must be explained by a small number of active features. Remarkably, the feature vectors obtained by fitting this model to natural images resemble the localized, oriented receptive fields found the early visual cortex (Olshausen & Field, 1996a). Subsequent work showed the model could account for a variety of properties of neural activity in the visual pathway (e.g. classical and non-classical receptive field effects (Rozell et al., 2008; Karklin & Lewicki, 2009; Lee et al., 2007)).

Although the sparse coding model is a linear generative model, simultaneous recognition (inferring the latent variables from an image) and learning (optimizing the dictionary of features) can be computationally intensive. Instead optimization has thus far relied on either variational optimization with a Dirac delta approximate posterior (Olshausen & Field, 1996a; Dayan et al., 2003; Seeger, 2008), which does not include uncertainty information, or sampling-based approaches (Berkes et al., 2008; Theis et al., 2012), which lack a neurally-plausible implementation. In fact, finding neurally-plausible architectures for both recognition and learning can be challenging in general. For variational methods, such architectures exist both for recurrent (Rozell et al., 2008; Charles et al., 2012; Zylberberg et al., 2011; Zhu & Rozell, 2013) and feed-forward (Gregor & LeCun, 2010), however these architectures rely on posterior approximations that do not include uncertainty.

In this paper, we propose a unified solution to these two important problems using ideas from the variational auto-encoder (VAE) (Kingma & Welling, 2014; Rezende et al., 2014). The VAE is a framework for training a complex generative model by coupling it to a recognition model parametrized by a deep neural network. This deep network offers tractable inference for latent variables from data and allows for gradient-based learning of the generative model parameters using a variational objective. Here we adapt the VAE methodology to the sparse coding model by adjusting its structure and prior assumptions. We compare the resulting *sparse-coding VAE* (SVAE) to fits using the original methodology, and show that our model achieves higher log-likelihood on test data. Furthermore, we show that the recognition model of the trained SVAE performs accurate inference under the sparse coding model, and captures important response properties of neurons in visual cortex, including orientation tuning, surround suppression, and frequency tuning.

## 2 Background

### 2.1 The Sparse Coding Model

The sparse coding model (Olshausen & Field, 1996a,b) posits that the spike response **z** from a population of neurons can be interpreted as a set of sparse latent variables underlying an image **x** presented to the visual system. This can be formalized by the following generative model:

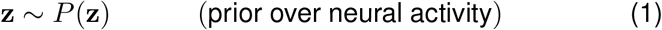

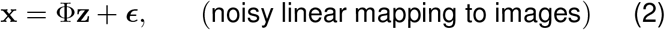

where *P*(**z**) is a sparse or heavy-tailed prior over neural activities, Φ is matrix of dictionary elements or “features”, and 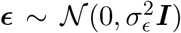 is isotropic Gaussian noise with variance 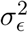. Critically, the sparse coding model is *overcomplete*, meaning that the dimension of the latent variable is larger than that of the data: Dim(**z**) *>* Dim(**x**). As a result, multiple settings of **z** can reproduce any given image patch **x**.

Although this linear generative model is easy to sample, the sparse prior over **z** make both inference and model fitting difficult. Full Bayesian inference involves computing the posterior distribution over latent variables given an image, which is (according to Bayes’ rule):

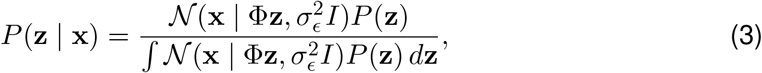

where 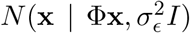 denotes a multivariate normal density in **x** with mean Φ **z** and covariance 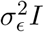. Unfortunately, the denominator cannot be computed in closed form for most sparsity-promoting priors, such as the Cauchy and Laplace distribution.

Olshausen & Field (1996a) considered a simpler inference problem by setting neural activity equal to the MAP estimate of **z** given the image:

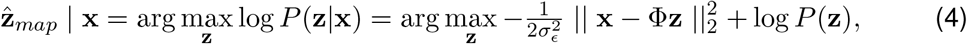

which does not depend on the denominator in (eq. 3). Here 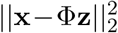 is the sum of squared errors between the image **x** and its reconstruction in the basis defined by Φ, and log *P*(**z**) serves as a “sparsity penalty” given by 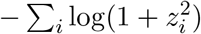 for a Cauchy prior or −Σ_*i*_ |*z*_*i*_| for a Laplace prior. The noise variance 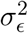 serves to trade off the importance of minimizing the image reconstruction error against the importance of sparsity imposed by the prior. (In the limit *σ*_*ϵ*_ → 0, the effect of the sparsity penalty log *P*(**z**) vanishes, and 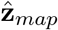 becomes a least-squares estimate). However, finding 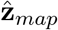 requires numerical optimization of the right-hand-side of (eq. 4), which does not easily map onto a model of feed-forward processing in the visual pathway.

### 2.2 Fitting the sparse coding model

The procedure for fitting the sparse coding model to data incurs additional steps beyond the inference problem. The maximum likelihood estimate of the model parameters Φ for a dataset of images *X* = {**x**_1_, …, **x**_*L*_} is given by

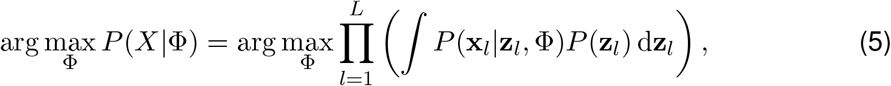

where *Z* = {**z**_1_, … **z**_*L*_} are the latent variables corresponding to the images in *X*. Once again, the integrals over **z**_*l*_ have no closed-form solution for the relevant sparsity-promoting priors of interest.

To circumvent this intractable integral, Olshausen & Field (1996a) employed an approximate iterative method for optimizing the dictionary Φ . After the initializing the dictionary randomly, iterate:

1. Take a group of training images *X*^(*i*)^ and compute the MAP estimate of the latent variables 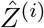 for each image using the current dictionary Φ^(*i*)^:

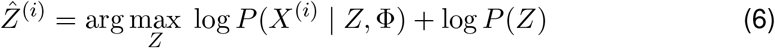
2. Update the dictionary using the gradient of the log-likelihood, conditioned on *Z*^(*i*)^:

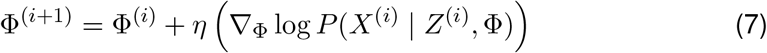

where *η* is a learning rate.

### 2.3 Fitting as Variational EM

The iterative fitting algorithm from (Olshausen & Field, 1996a,b) can be seen to relate closely to a form variational expectation-maximization (EM) (Olshausen, 1996; Dayan et al., 2003). Variational EM employs a surrogate distribution *Q*(*Z*) to optimize a tractable lower-bound on the log-likelihood known as the *evidence lower bound* (ELBO):

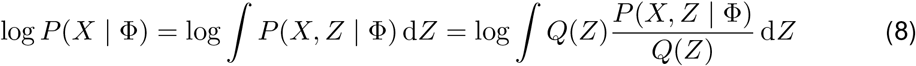

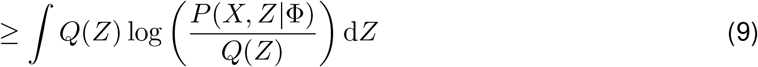

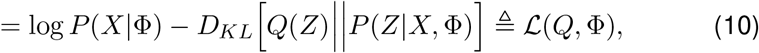

where the inequality in (eq. 9) follows from Jensen’s inequality, and *D*_*KL*_[*Q*(*Z*)||*P*(*Z*)] =∫ *Q*(*Z*) log[*Q*(*Z*)*/P* (*Z*)]*dZ* denotes the Kullback-Leibler (KL) divergence between two distributions *Q*(*Z*) and *P*(*Z*) (Bishop, 2005; Blei et al., 2016).

The expectation or “E” step of variational EM involves setting *Q*(*Z*) to minimize the KL-divergence between *Q*(*Z*) and *P*(*Z*|*X*, Φ ), the posterior over the latent given the data and model parameters. Note that when *Q*(*Z*) = *P*(*Z*|*X*, Φ ), the KL term (Eq. 10) is zero and the lower bound is tight, so *ℒ* achieves equality with the log-likelihood. The maximization or “M” step involves maximizing *ℒ*(*Q*, Φ ) for Φ with *Q* held fixed, which is tractable for appropriate choices of surrogate distribution *Q*(*Z*). Variational EM can therefore be viewed as alternating coordinate ascent of the ELBO:

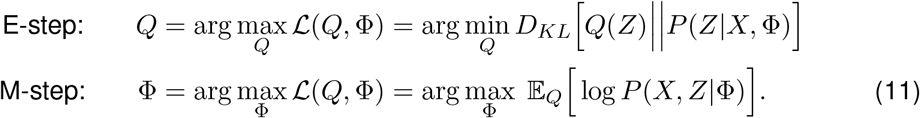

It is now possible to see that the fitting algorithm of Olshausen & Field (1996a), as also discussed by Dayan et al. (2003) is closely related to variational EM in which the surrogate distribution *Q*(*Z*) is a product Dirac delta functions:

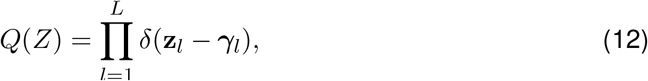

where {***γ***_1_, …, ***γ***_*L*_} are the variational parameters governing *Q*. With a Dirac delta posterior, the EM algorithm is formally undefined. Examining Equation 11, the KL divergence in the E-step between the Dirac delta distribution and the latent variable posterior is infinite, and the ELBO is formally −∞, always. We can see the source of this catastrophe by writing out the KL divergence of the E-step above as

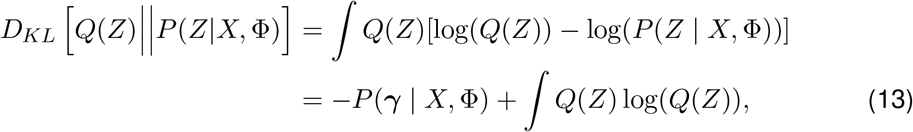

where in the second line we have used our delta-function variational distribution *Q*(*X*) (eq.12) to evaluate the first term, which is simply the posterior evaluated at ***γ***. The second term is infinite; note however that this infinite term is independent of Φ, and so the gradient of the KL divergence with respect to Φ can be computed sensibly, and optimized to perform the E-step.

If we neglect the infinite term (which is constant in Φ ), the ELBO reduces to:

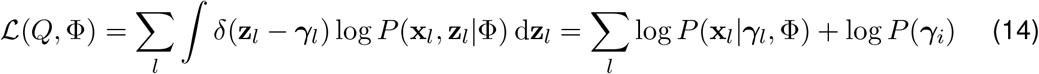

The E-step involves setting ***γ***_*l*_ to the MAP estimate of the latent vector **z**_*l*_ for each image **x**_*l*_, while the M-step involves updating the dictionary Φ to minimize the reconstruction error of all images given the inferred latent variables. The original sparse coding algorithm differed in that it operated on randomly selected sub-sets of the data (stochastic mini-batches), and took a single gradient step in Φ in place of a full M-step at each iteration. The algorithm can therefore be seen as a stochastic gradient descent version of variational EM (R. Neal & Hinton, 1998).

Employing a Dirac delta variational posterior introduces bias whereby Φ can grow without bound to increase the ELBO^1^. In the case that a variational distribution with finite variance is used, the norm of Φ is regularized by the KL divergence. To compensate for this bias, Olshausen and Field introduced a post-hoc normalization procedure which effectively matches the **z** variance to the data variance, regularizing Φ in the process. Interestingly, we can also show how this step can arise in the variational framework. We first write the M-step, ignoring terms not depending on Φ, and assume for now that *Q*(*Z*) has mean **z**_*map*_ and covariance Σ_*z*_:

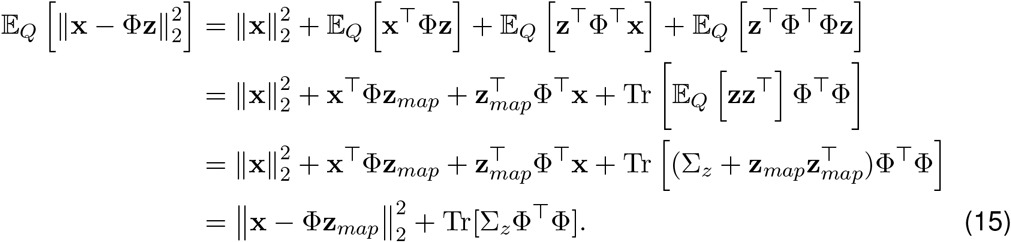

If all datapoints were accessible at once, we can solve this optimization directly. In the case of random mini-batches, we must take small gradient steps. Specifically we consider the proximal gradient descent algorithm (Boyd & Vandenberghe, 2004) for optimizing over a batch of data points *X* = [**x**_1_, **x**_1_, …],

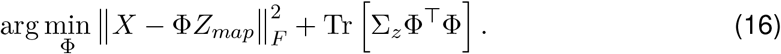

In proximal gradient descent, optimizing a general cost *J*(Φ ) = *f* (Φ ) + *g*(Φ ), where *f* (*·*) is smooth and *g*(*·*) is potentially non-smooth, can be solved by alternating between gradient steps with respect to *f* (*·*),

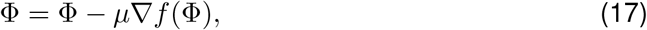

and updating the solution via a proximal projection

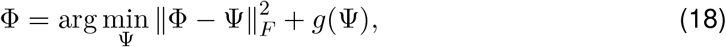

which can often be solved in closed form. For example, the proximal gradient algorithm for the LASSO estimator reduces to the well known iterative soft-thresholding algorithm (ISTA) (Beck & Teboulle, 2009).

While *g*(*·*) here is trivially smooth (i.e., a quadratic cost), we can still apply proximal gradient descent, obtaining the following updates:

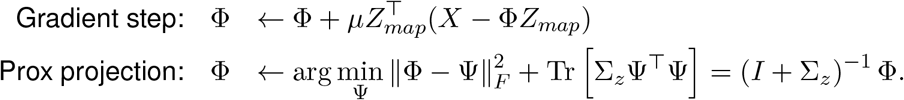

A slight re-organization in the projection step gives

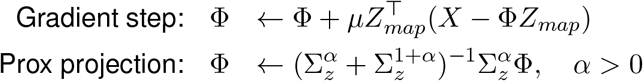

which for diagonal Σ_*z*_ introduces a per-dictionary normalization

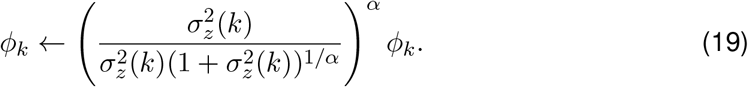

Note that this update step has the same form as the re-normalization step in the original sparse coding algorithm, with 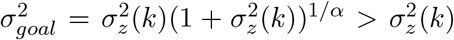. The strict inequality here ensures that the variance of Φ is larger than the variance in the latents, which is what drives the solution away from the trivial solution. The degree of how much greater it is is dictated by the parameter *α*. One way, then, to interpret this algorithm then is to assume that in this step, Σ_*z*_ is replaced by the empirical covariance across samples before the limit as Σ_*z*_ → 0 for the Dirac-delta approximation. Interestingly, in the case where a prior over Φ is used where 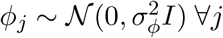, the same process holds, except the proximal update is

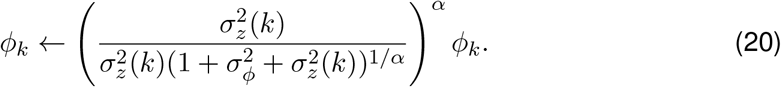

where now 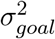 can be modulated arbitrarily in the range 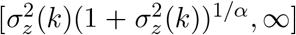. This is interesting since it indicates that the update step in the original sparse coding is implicitly implementing the regularization only introduced explicitly in later work (Karklin & Lewicki, 2005).

### 2.4 Variational Auto-Encoders (VAEs)

The variational auto-encoder is a powerful framework for training complex generative models (Kingma & Welling, 2014; Rezende et al., 2014; Doersch, 2016). In the first papers describing the VAE, the generative model was defined by a standard Gaussian latent variable mapped through a deep neural network with additive Gaussian noise:

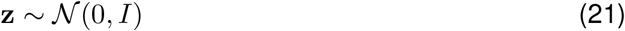

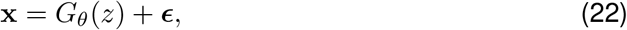

where *G*_*θ*_(**z**) denotes the output of a deep neural network with parameters *θ*, and 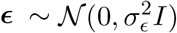 represents Gaussian noise added to the output of this network. By using a sufficiently large, deep network, this model should be able to generate data with arbitrarily complicated continuous distributions.

The key idea of the VAE framework is to perform variational inference for this model using a variational distribution parametrized by another deep neural network:

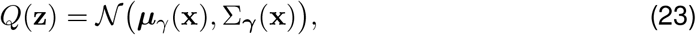

where ***μ***_*γ*_(**x**) and Σ_***γ***_(**x**) denote the neural network with parameters ***γ*** = {*γ*_*μ*_, *γ*_*σ*_}, that map data samples **x** to the mean and covariance (respectively) of a Gaussian variational distribution over **z** given **x**. These network outputs provide an approximate recognition model for inferring **z** from **x** under the generative model (Eqs. 21-22). In VAE parlance, ***μ***_*γ*_(*·*) and Σ_***γ***_(*·*) comprise the *encoder*, mapping data samples to distributions over the latent variables, while the generative network *G*_*θ*_(*·*) is the *decoder*, mapping latent variables to the data space. This terminology is inspired by the model’s structural similarity to classic auto-encoders, and is consistent with the fact that **z** is typically lower-dimensional than **x**, providing a compressed representation of the structure contained in the data when using the encoder. Note that the analogy is imprecise, however: the output of the encoder in the VAE is not a point **z** in latent space, but a distribution over latent variables *Q*(**z**).

Fitting the VAE to data involves stochastic gradient ascent of the ELBO (Eq. 10) for model parameters *θ* and variational parameters ***γ***. This can be made practical using several clever tricks. The first trick is to evaluate the expectation over *Q*(*Z*) as a Monte Carlo integral. The contribution to the ELBO from a single data sample **x** is therefore:

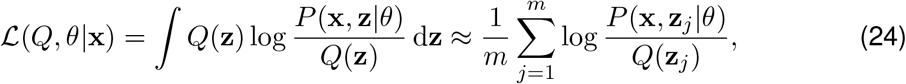

for **z**_1_, … **z**_*m*_ ∼ *Q*(**z**), where *m* is the number of samples used to evaluate the Monte Carlo integral.

The second trick is the *reparametrization trick*, which facilitates taking derivatives of the above expression with respect to the variational parameters ***γ*** governing *Q*(**z**). Instead of sampling **z**_*j*_ ∼ *Q*(**z**), the idea is to map samples from a standard normal distribution into samples from the desired variational distribution *Q*(**z**) = 𝒩 (***μ***_*γ*_(**x**), Σ_***γ***_(**x**)) via a differentiable transformation, namely:

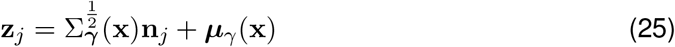

for **n**_1_, …, **n**_*m*_ ∼ *N* (0, *I*), where 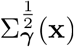 denotes the matrix square root of the covariance matrix Σ_***γ***_(**x**).

Combining these two tricks, and plugging in the VAE generative and recognition model components for *P*(**x, z**|*θ*) and *Q*(**z**), a Monte Carlo evaluation of the per-datum ELBO can be written:

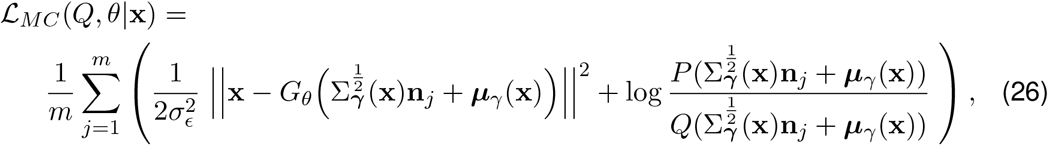

where the first term is the the mean squared error between the original image patch **x** and its reconstruction after passing **x** through the noisy encoder and decoder, and the second term is the log-ratio of the latent prior *P*(**z**) = 𝒩 (0, *I*) and the variational distribution *Q*(**z**) = *N* (***μ***_*γ*_(**x**), Σ_***γ***_(**x**)), evaluated at the sample point 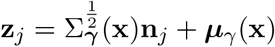.

It is worth noting that the first term in (eq. 26) relies on a stochastic pass through the encoder, given by a noisy sample of the latent from *Q*(**z**) (which is really an approximation to *P*(**z**|**x**), the conditional distribution of the latent given **x**). This sample is then passed deterministically through the decoder network *G*_*θ*_. The generative model noise variance 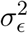 serves as an inverse weight that determines how much to penalize reconstruction error relative to the second term in the ELBO. The second term, in turn, can be seen as a Monte Carlo estimate for −*D*_*KL*_(*Q*(**z**)||*P*(**z**)), the negative KL divergence between the variational posterior and the prior over **z**. Because both these distributions are Gaussian, the standard approach is to replace the Monte Carlo evaluation of this term with its true expectation, using the fact that the KL divergence between two Gaussians can be computed analytically (Kingma & Welling, 2014; Rezende et al., 2014).

In contrast to the iterative variational EM algorithm used for the classic sparse coding model, optimization of the VAE is carried out by simultaneous gradient ascent of the ELBO with respect to generative network parameters *θ* and variational parameters ***γ***. During training, the per-datum ELBO in (eq. 26) is summed over a mini-batch of data for each stochastic gradient ascent step.

## 3 Sparse Coding VAEs

In this paper, we adapt the VAE framework to the sparse coding model, a sparse generative model motivated by theoretical coding principles. This involves three changes to the standard VAE: (1) We replace the deep neural network from the VAE generative model with a linear feed-forward network; (2) We change the under-complete latent variable representation to an overcomplete one, so that the dimension of the latent variables is larger than the dimension of the data; and (3) We replace the standard normal prior over the latent variables with heavy-tailed, sparsity-promoting priors (e.g., Laplace and Cauchy).

We leave the remainder of the VAE framework intact, including the conditionally Gaussian variational distribution *Q*(**z**), parametrized by a pair of neural networks that output the mean and covariance as a function of **x**. The parameters of the SVAE are therefore given by 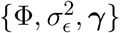and a prior *P*(**z**), where *P*(**z**), Φ and 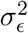 specify the elements of sparse coding model (a sparse prior, generative weight matrix, and Gaussian noise variance, respectively), and variational parameters ***γ*** are the weights of the recognition networks ***μ***_*γ*_(*·*) and Σ_***γ***_(*·*) governing the variational distribution *Q*(**z**) = 𝒩 (*μ*(**x**), Σ_***γ***_(**x**)). Fig. 2 shows a schematic comparing the two models.

**Figure 1:**
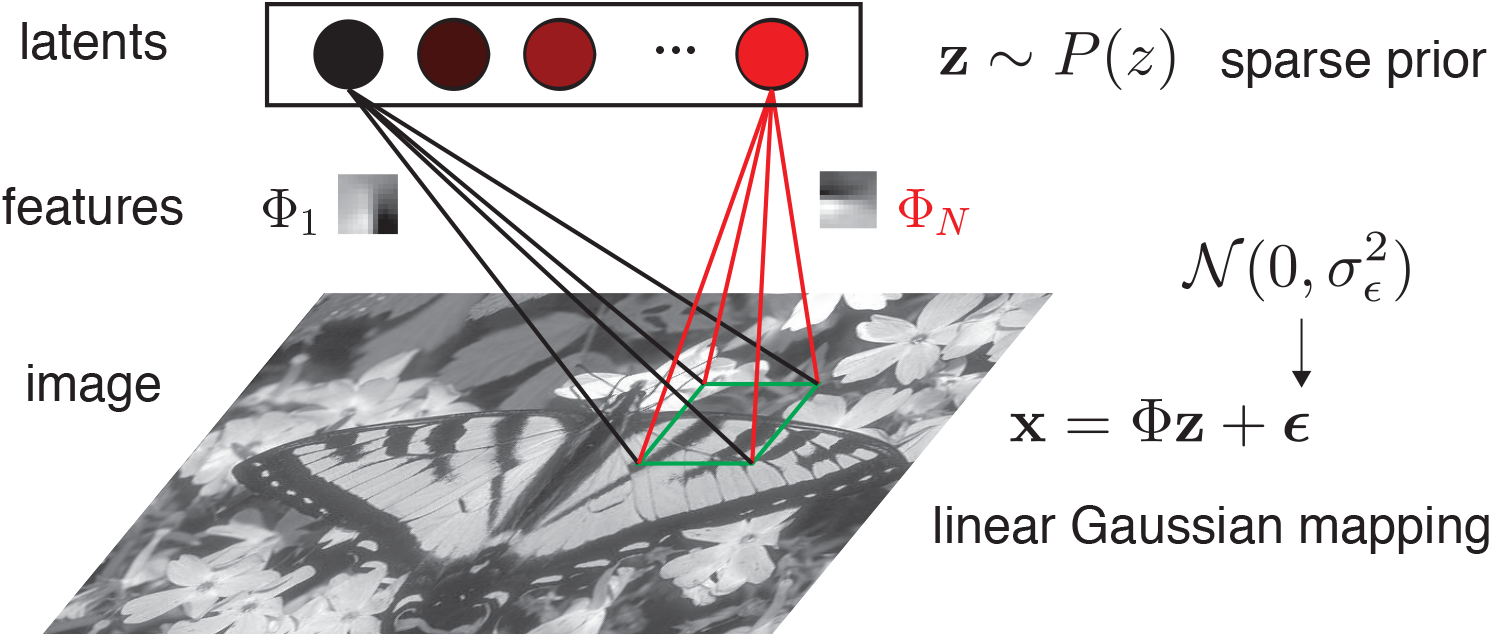
The sparse coding model. A sample from the model is generated by first sampling the latent variables **z** from the sparse prior distribution *P*(**z**), which provide linear weights on feature vectors Φ_*i*_, and finally adding Gaussian noise. The image shown is taken from the BSDS300 dataset (Martin et al., 2001).

**Figure 2:**
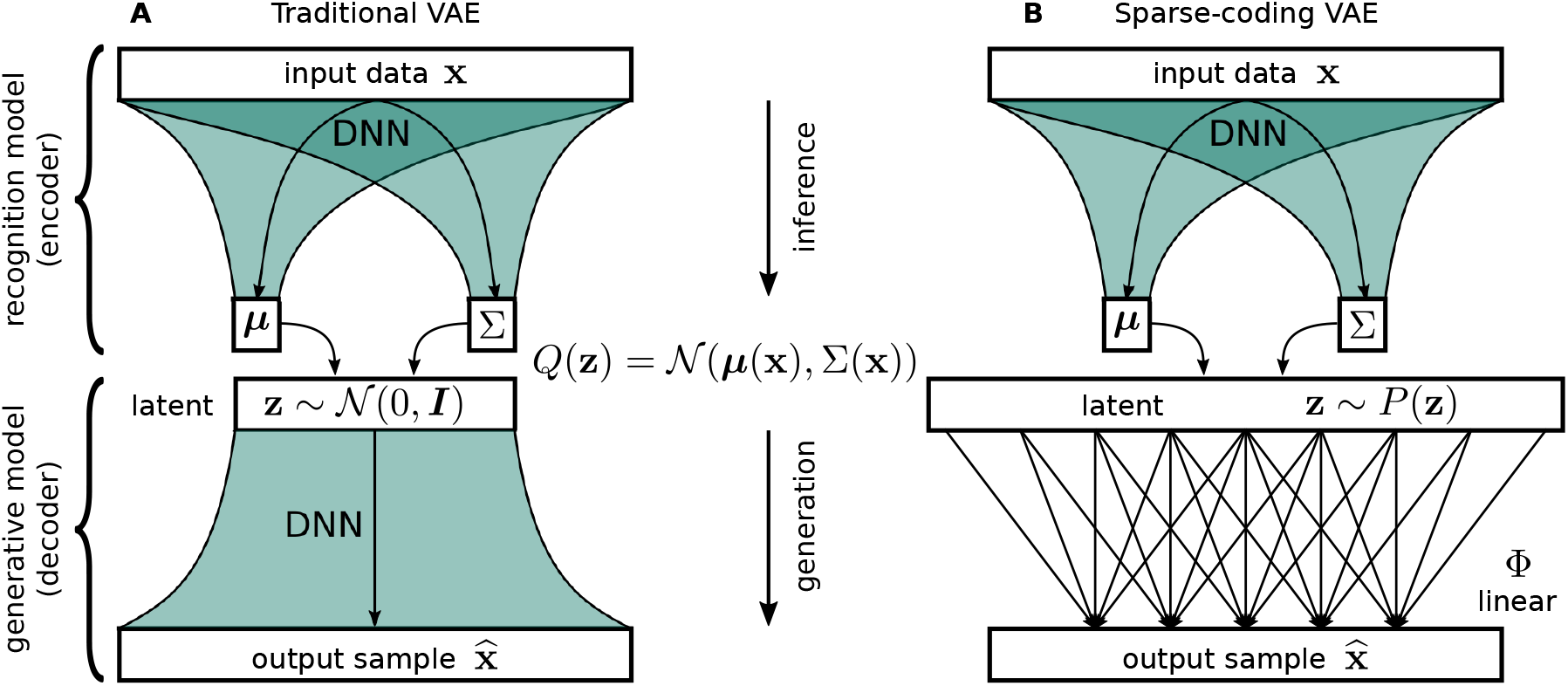
Schematics of traditional VAE and the sparse-coding VAE. **(A)** The traditional VAE consists of a generative model (decoder) with a recognition model (encoder) stacked on top to subserve variational inference. Both models are parametrized by deep neural networks with Gaussian noise. **(B)** The sparse-coding VAE results from replacing the generative model by the sparse coding model. It differs from the original VAE in three key respects: the latent variable **z** has higher dimension than the data **x**; the prior over **z** is sparse (e.g., Cauchy or Laplace) instead of Gaussian; and the generative model is linear instead of a deep neural network.

## 4 Methods

### 4.1 Data Preprocessing

We fit the SVAE to 12 *×* 12 pixel image patches sampled from the BSDS300 dataset (Martin et al., 2001). Before fitting, we preprocessed the images and split the data into train, test, and validation sets. In the original sparse coding model, the images were whitened in the frequency domain, followed by a low-pass filtering stage. In Olshausen & Field (1996a) the whitening step was taken to expedite learning, accentuating high frequency features that would be far less prominent for natural image data, which is dominated by the low-frequency features. This is due to the fact that the Fourier (frequency) components are approximately the principal components of natural images, however the overall variance of each component scales with the inverse frequency squared (the well known 1*/f* spectral properties of natural images), producing large differences in variance between the high- and low-frequency features. This poses a problem for gradient-based approaches to fitting the sparse coding model since low variance directions dominate the gradients. The low-pass filtering stage then served to reduce noise and artifacts from the rectangular sampling grid.

We perform a slight variation of these original preprocessing steps, but with the same overall effect. We whiten the data by performing PCA on the natural images and normalizing each component by its associated eigenvalue. Our low-pass filtering is achieved by retaining only the 100*π/*4% most significant components, which correspond to the 100*π/*4% lowest frequency modes, which also roughly corresponds to a circumscribed circle in the square Fourier space of the image, removing the noisy, high frequency corners of Fourier space.

### 4.2 SVAE Parameters

We implemented an SVAE with dim(**z**) = 169 latent dimensions, resulting in a latent code that is 1.5*×* over-complete, and set the Gaussian output noise variance to 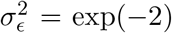 (implying a signal-to-noise ratio of exp(2), since the data is whitened). We also tested a latent code that is 2*×* over-complete and observed similar results, and thus kept *×*1.5 over-completeness for computational considerations. We implemented the SVAE with three choices of prior which included Laplace (Eq. 27), Cauchy (Eq. 28), and Gaussian (Eq. 29).

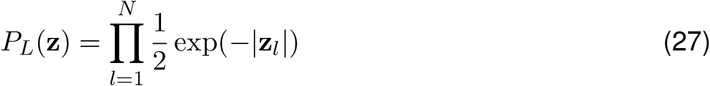

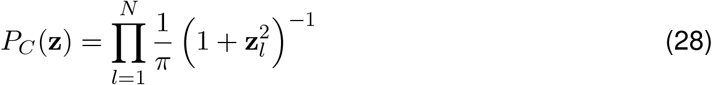

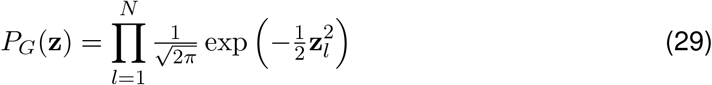

We parametrized the recognition models ***μ***_*γ*_(*·*) and Σ_***γ***_(*·*) using feed-forward deep neural networks (Figure 2). The two networks took as input a vector of preprocessed data and had a shared initial hidden layer of 128 rectified linear units. Each network had two additional hidden layers of 256 and 512 rectified linear units respectively, which were not shared. These networks, parameterizing ***μ***_*γ*_(*·*) and Σ_***γ***_(*·*), each output a dim(**z**) = 169 dimensional vector which encode the mean and main diagonal of the covariance of the posterior, respectively. We set the off-diagonal elements of the posterior covariance matrix to zero — this assumption, and its implications for the dependencies between latent coefficients, is further discussed in the Discussion section. The final hidden layer in each network is densely connected to the output layer, with no nonlinearity for the mean output layer and a sigmoid nonlinearity for the variance output layer. In principle the variance values should be encoded by a non-saturating positive-definite nonlinearity; however we found that this led to instability during the fitting process and the sigmoid nonlinearity resulted in more stable behavior. Intuitively, given that our priors have scales of one, the posteriors will generally have variances less than 1, and can be expressed sufficiently well with the sigmoid nonlinearity.

### 4.3 Optimization

We optimized the SVAE using the `PyTorch` (Paszke et al., 2017) machine learning framework. Gradient descent was performed for 128 epochs (approximately 2 *×* 10^5^ optimization steps) with the Adam optimizer (Kingma & Ba, 2014) with default parameters, a batch size of 32, and a learning rate of *η* = 10^−4^ . The networks always converged well within this number of gradient descent steps. We took the number of Monte-Carlo integration samples to be *m* = 1 and tested higher values, up to *m* = 5, but found that this parameter did not influence the results. We used the same learning hyperparameters for all three priors.

For comparison we fit our own version of the sparse coding model, also cast as a PyTorch “Module”, using the methods of Olshausen and Field (Olshausen & Field, 1996a) for fitting . We utilized the same learning hyperparameters as in the original work, including during normalization of Φ (see Section 2.3).

### 4.4 Evaluating Goodness of Fit with Annealed Importance Sampling

To evaluate the goodness of fit we used annealed importance sampling (AIS) (R. M. Neal, 2001). We refer to Appendix §8.1 for a primer on the topic, and the use of the method for evaluating log-likelihoods. Our estimates used 1000 samples with 16 independent chains, a linear annealing procedure over 200 intermediate distributions, and a transition operator consisting of one HMC trajectory with 10 leapfrogs steps. Furthermore, we tuned the HMC step-size to achieve (within 1% absolute tolerace) the optimal acceptance rate of 0.65 (R. M. Neal, 2011; Wu et al., 2016). We do so using a simple gradient descent algorithm: consecutively over single batches of input samples, denoting *A*_*ϵ*_ the average HMC acceptance rate obtained when computing AIS on those input samples, update *ϵ* ← *ϵ* + *η* (*A*_*ϵ*_ − 0.65) with learning rate *η* ≃ 0.5 until convergence. Finally, we note that AIS only gives a lower bound on the log-likelihood (Wu et al., 2016). Even though an exact expression is available for the log likelihood in the Gaussian case, we use AIS for the Gaussian prior as well so that the resulting values are directly comparable.

## 5 Results

### 5.1 Quality of Fit

We first assess our model by calculating goodness-of-fit measures. We compare here the various choices in prior distributions using both the SVAE optimized with VAE-based inference and the original sparse coding model fit with the method of (Olshausen & Field, 1996a). To perform this assessment, we computed the log-likelihood of test data under the fit parameters, using AIS. Figure 3A depicts how the log-likelihood is monotonically increasing over training for all priors. We observe that, of the priors tested, sparser distributions result in higher log-likelihoods, with the Cauchy prior providing the best fit. To explore the utility of the sparse coding VAE over the approximate EM method, we similarly calculated the log-likelihoods for the method in (Olshausen & Field, 1996a), again for each of the three prior distributions. We observe that the log-likelihood goodness-of-fit for the VAE is higher than the equivalent fits obtained with the method of (Olshausen & Field, 1996a) (Table 1). This is due to the fact that the sparse coding VAE uses a more robust approximation of the log-likelihood, and the variational posterior of VAEs are more informative than simply using the posterior mode. Indeed, we report in Figure 3B the inferred latent values for this variational posterior in comparison to the posterior mode. The closer the approximation is to the *true* posterior, the closer this histogram of latent values would be to the prior. We see that the inferred latent values of the sparse coding VAEs provide a better approximation of the prior than those obtained with the Dirac-delta approximation.

**Table 1:** Log-likelihood calculations for the sparse coding VAE (top) versus the traditional Dirac-delta approximation. Values were calculated using AIS with HMC transitions, and show improvement across all prior choices.

| (nats)<br>Implementation | Prior |  |  |
| --- | --- | --- | --- |
|  | Cauchy | Laplace | Gaussian |
| VAE | **57** | **50** | **27** |
| Traditional method | -221 | -117 | -127 |

**Figure 3:**
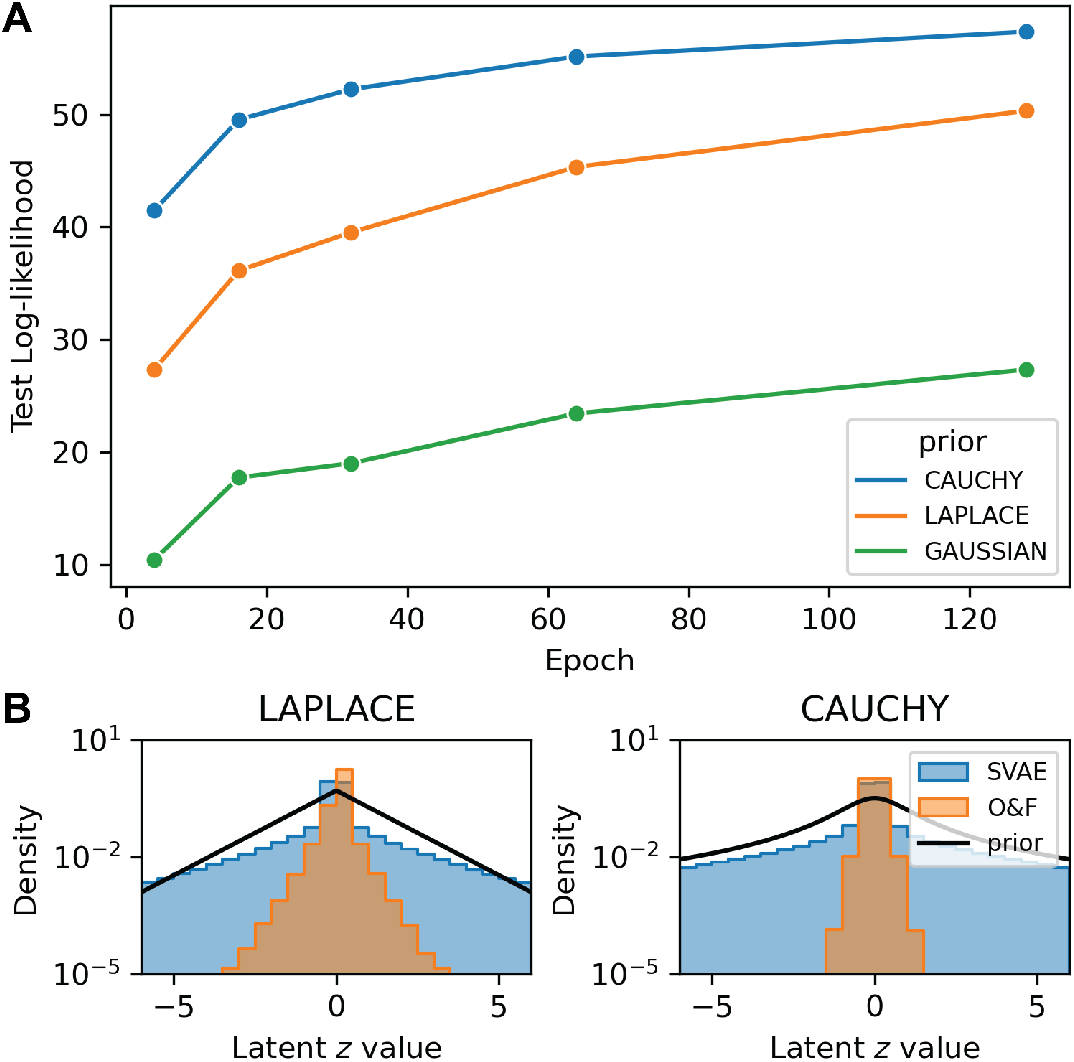
Quantitative analysis of quality of fit. **Panel A**: Log-Likelihood of VAE fit throughout training for Laplace, Cauchy and Gaussian priors. **Panel B**: Histogram of inferred latent variables sampled (10K samples)from the variational posterior under our fit and the fit method of (Olshausen & Field, 1996a) (“O&F” label) along with the true prior (black line), for both Laplace (left) and Cauchy (right) priors . Note that the variational posterior in the method of (Olshausen & Field, 1996a) is a Dirac-delta distribution at the MAP estimate of **z**.

Finally, we compare in Figure 4 how the learned basis functions in the sparse coding VAE framework differ from those learned in the standard sparse coding (Olshausen & Field, 1996a). We analyzed the learned basis functions by fitting a Gabor filter to each feature, and report the estimated frequency and orientation distributions (see Appendix 8.2 for more details). We observe a persistent increase in frequency across all priors for the SVAE over the standard sparse coding. For the orientation, there is less of a discernible trend, but we do generally observe more variable orientation distribution for the SVAE basis functions, as opposed to the more flat distribution for the standard sparse coding.

**Figure 4:**
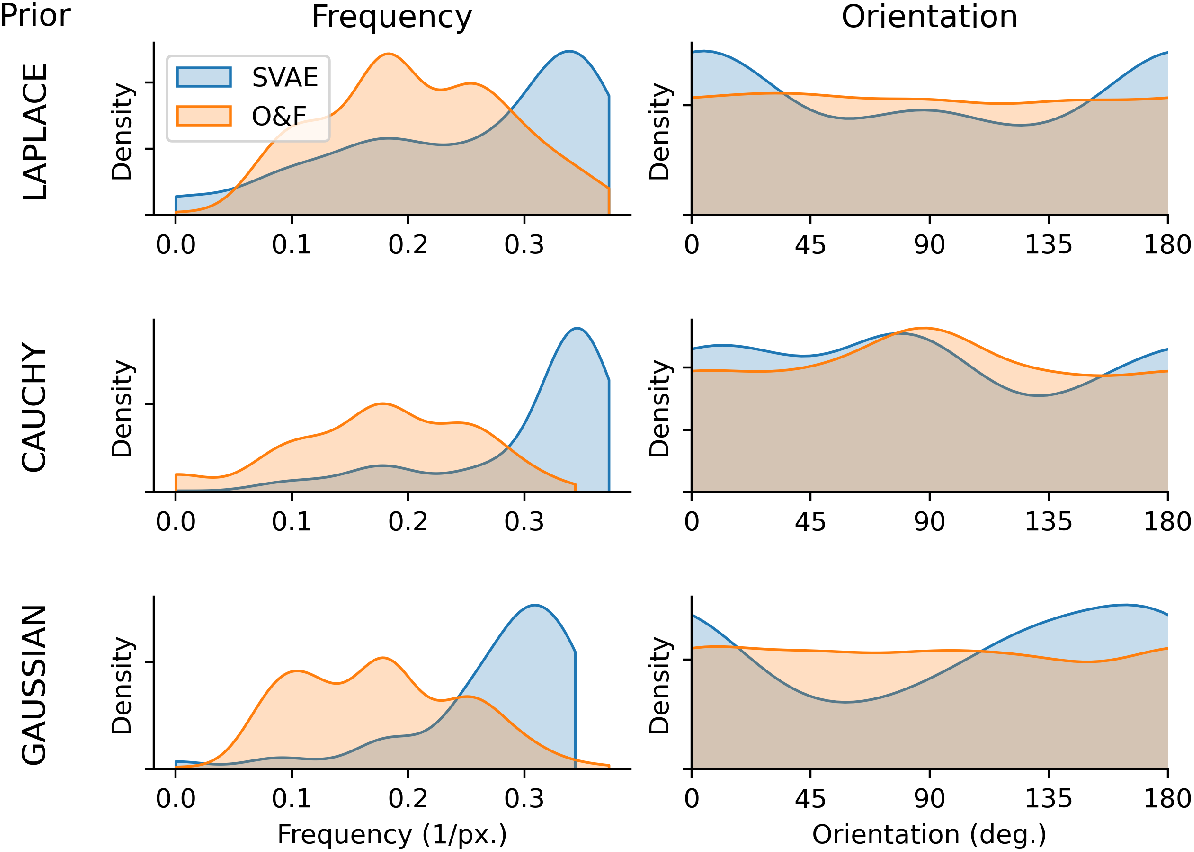
Frequency and orientation statistics for the Gabor filter fits to the learned basis functions. For each basis function we obtain a frequency and orientation statistic, and we report the Kernel density estimate (KDE) of the distribution. The rows indicate which prior was used over the latent coefficients, and the color in each plot indicates the inference methodology (ours against the original sparse coding from Olshausen & Field (1996a) (“O&F”))

### 5.2 Feed Forward Inference Model

As a consequence of training a VAE, we obtain a neural network which performs approximate Bayesian inference. Previous mechanistic implementations of sparse coding use the MAP estimates under the true posterior to model trial-averaged neural responses (Rozell etal., 2008; Boerlin & Denève, 2011; Gregor & LeCun, 2010; Martins et al., 2011). In our case, the recognition model performs approximate inference using a more expressive approximation, suggesting that it may serve as an effective model of visual cortical responses. To study the response properties of the feed-forward inference network we simulated neural activity as the mean of the recognition distribution *Q*(**z**) ≈ *P*(**z**|**x**) with stimulus **x** taken to be sinusoidal gratings of various sizes, contrasts, angles, frequencies, and phases. For each set of grating parameters, we measured responses to both a cosine and sine phase grating. To enforce non-negative responses and approximate phase invariance, the responses shown in Figure 5 are the root sum-of-squares of responses to both phases.

**Figure 5:**
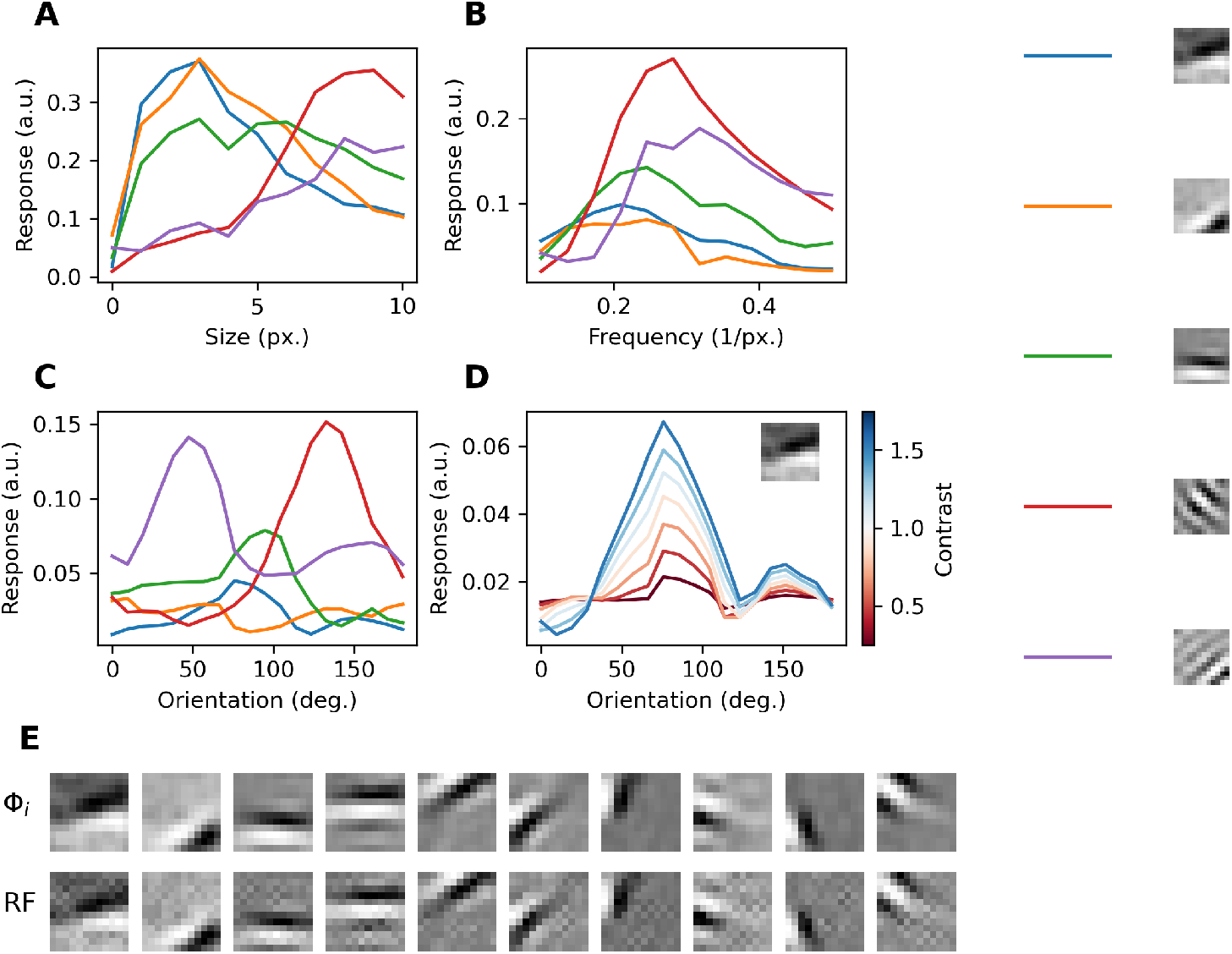
Visualization of feed-forward inference model behavior. Curves in Panels A-C are generated using full-contrast gratings. Dictionary elements associated with each neuron are visualized on the right. **Panel A**: size tuning curves of five sample neurons in response to a grating with optimal orientation and frequency. **Panel B**: Frequency response function of five model neurons in response to a full-field, full-contrast grating with optimal orientation. **Panel C**: Orientation preference curves of five model neurons in response to full-field, full-contrast gratings. **Panel D**: Example orientation preference curves of a single model neuron in response to a full-field grating at varying contrasts (denoted by line darkness) demonstrating contrast invariance. **Panel E**: Comparison of dictionary elements (top) with receptive fields (bottom) measured as in a reverse-correlation experiment for measuring receptive field (Ringach & Shapley, 2004). The stimulus used for reconstruction was Gaussian white noise and receptive fields were averaged over 10000 random stimulus presentations.

Fig. 5 shows the performance of the recognition network in response to sinusoidal gratings of various sizes, contrasts, angles, and phases. We found that the responses of the recognition model exhibited frequency and orientation tuning (Fig. 5B,C), reproducing important characteristics of cortical neurons (Hubel & Wiesel, 1962) and reflecting the Gabor-like structure of the dictionary elements. Additionally, Fig. 5D demonstrates that the orientation tuning of these responses is invariant to grating contrast, which is observed in cortical neurons (Troyer et al., 1998). Fig. 5A shows that the recognition model exhibits the surround suppression, in which the response to an optimally-tuned grating first increases with grating size and then decreases as the scale of the grating increases beyond the classical receptive field (Sceniak et al., 1999). Lastly, Figure 5E shows the receptive fields of recognition model neurons, as measured by a reverse-correlation experiment (Ringach & Shapley, 2004), verifying that the linear receptive fields exhibit the same Gabor-like properties as the dictionary elements.

## 6 Discussion

We have introduced a variational inference framework for the sparse coding model based on the VAE. The resulting SVAE model offers a more principled and accurate method for fitting the sparse coding model, and comes equipped with a neural implementation of feed-forward inference under the model. We showed, first of all, that the classic fitting method of Olshausen & Field is equivalent to variational inference under a delta-function variational posterior. We then extended the VAE framework to incorporate the sparse coding model as generative model. In particular, we replaced the standard deep network of the VAE with an overcomplete latent variable governed by a sparse prior, and showed that variational inference using a conditionally Gaussian recognition distribution provided accurate, neurally plausible feed-forward inference of latent variables from images. Additionally, the SVAE provided improved fitting of the sparse coding model to natural images, as measured by the test log-likelihood. Moreover, we showed that the associated recognition model recapitulates important response properties of neurons in the early mammalian visual pathway.

Given this demonstration of VAEs for fitting tailor-made generative models, it is important to ask whether VAEs have additional applications in theoretical neuroscience. Specifically, many models are constrained by their ability to be fit. Our technique may allow more powerful, yet still highly structured, generative models to be practically applied by making learning tractable. The particular property that VAEs provide an explicit model of the posterior (i.e. as opposed to the Dirac delta approximation) means that connections can also now be drawn between generative models, (e.g., sparse coding) and models that depend explicitly on neural variability — a property often tied to the confidence levels in encoding. Current models of over-dispersion (e.g. (Goris et al., 2014; Charles et al., 2018)) shy from proposing mechanistic explanations of the explanatory statistical models. Another important line of future work then is to explore whether the posterior predictions made by SVAEs, or VAEs tuned to other neuroscience models, can account for observed spiking behaviors.

### Relationship to previous work

The success of the sparse coding model as an unsupervised learning method for the statistics of the natural world has prompted an entire field of study into models for sparse representation learning and implementations of such models in artificial and biological neural systems. In terms of the basic mathematical model, many refinements and expansions have been proposed to better capture the statistics of natural images, including methods to more strictly induce sparsity (Garrigues & Olshausen, 2008; Girolami, 2001; Olshausen & Millman, 2000; M. S. Lewicki & Olshausen, 1999), hierarchical models to capture higher order statistics (Karklin & Lewicki, 2003, 2005, 2009; Garrigues & Olshausen, 2010), sampling-based learning techniques (Berkes et al., 2008; Theis et al., 2012) constrained dictionary learning for non-negative data (Charles et al., 2011), and applications to other modalities, such as depth (Tǒsic et al., 2011), motion (Cadieu & Olshausen, 2009) and auditory coding (Smith & Lewicki, 2006).

Given the success of such models in statistically describing visual responses, the mechanistic question as to how a neural substrate could implement sparse coding also became an important research area. Neural implementations of sparse coding have branched into two main directions: recurrent (Rozell et al., 2008; Boerlin & Denève, 2011) and feed-forward neural networks (Gregor & LeCun, 2010; Martins et al., 2011). The recurrent network models have been shown to provably solve the sparse-coding problem (Rozell et al., 2008; Shapero et al., 2014; Schwemmer et al., 2015). Furthermore, recurrent models can implement hierarchical extensions (Charles et al., 2012) as well as replicate key properties of visual cortical processing, such as non-classical receptive fields (Zhu & Rozell, 2013). The feed-forward models are typically based off of either mimicking the iterative processing of recursive algorithms, such as the iterative soft-thresholding algorithm (ISTA) (Daubechies et al., 2004) or by leveraging unsupervised techniques for learning deep neural networks, such as optimizing auto-encoders (Makhzani & Frey, 2013). The resulting methods, such as the learned ISTA (LISTA) (Gregor & LeCun, 2010; Borgerding et al., 2017), provide faster feed-forward inference than their RNN counterparts^2^, at the cost of losing theoretical guarantees on the estimates.

Although the details of these implementations vary, all essentially retain the Dirac-delta posterior approximation and are thus constructed to calculate MAP estimates of the coefficients for use in a gradient-based feedback to update the dictionary. None of these methods re-assess this basic assumption, and so are limited in the overall accuracy of the marginal-log-likelihood estimation of the model Φ, as well as their ability to generalize beyond inference-based networks to other theories of neural processing, such as probabilistic coding (Fiser et al., 2010; Orbán et al., 2016). In this work we have taken advantage of the refinement of VAEs in the machine learning literature to revisit this initial assumption from the influential early work and create just such a posterior-seeking neural network. Specifically, VAEs can provide a nontrivial approximation of the posterior via a fully Bayesian learning procedure in a feed-forward neural network model of inference under the sparse coding model. Additional benefits of VAEs over non-variational autoencoders with similar goals (e.g., LISTA) are the emerging properties of robustness to outliers and local minima, as observed in recent analysis (Dai et al., 2018). Recent work by Velychko et al. (2023) has also leveraged the formalism offered by the variational framework, focusing on analytical and entropy-based derivations of the ELBO instead of the parallelism with the original sparse coding work presented here. As VAEs have advanced, so have their abilities to account for more complex statistics in the latent representation layer. Discrete-type distributions, such as the enabled by the Concrete or Gumbal Softmax distributions enable categorical modeling that more closely resembles a version of sparsity (Van Den Oord et al., 2017; Maddison et al., 2016a; Jang et al., 2016). Nonlinear ICA (Khemakhem et al., 2020), or other disentangling methods (Chen et al., 2018) seek distributions that maximize independence between the latent representation variables, however rely on highly nonlinear decoding networks, removing interpretability as related to the data itself.

Obtaining the posterior distribution is especially important given that neural variability and spike-rate over-dispersion can be related to the uncertainty in the generative coefficients’ posterior (e.g., via probabilistic coding (Fiser et al., 2010; Orbán et al., 2016)). Finding a neural implementation of sparse coding that also estimates the full posterior would be an important step towards bridging the efficient and probabilistic coding theories. Other variational sparse coding models tended to sacrifice neurally-plausible implementation (Berkes et al., 2008; Theis et al., 2012; Seeger, 2008). Our work complements recent efforts to connect the sparse coding model with tractable variational inference networks. These works have focused on either the non-linear sparsity model (Salimans, 2016), or the linear generative model of sparse coding (Aitchison et al., 2018). Our work can also be thought of as generalizing models that use, for example Expectation Propagation (EP), to approximate the posterior distribution (Seeger, 2008). Other related work uses traditional VAEs and then performs sparse coding in the latent space (Sun et al., 2018). To date, however, no neurally-plausible variational method has been designed to capture the three fundamental characteristics of sparse coding: overcomplete codes, sparse priors, and a linear-generative model.

### Limitations and future directions

The state-of-the-art results of this work are primarily the result of orienting sparse coding in a variational framework where more expressive variational posterior distributions can be used for model fitting. Nevertheless this work represents only the first steps in this direction. One area for improvement is our selection of a Gaussian variational posterior with diagonal covariance matrix Σ_***γ***_(*·*). This choice was meant to expand the Dirac-delta posterior approximation of traditional sparse coding to include uncertainty. This model, however, still restricts the latent variables to be uncorrelated under the posterior distribution and can limit the variational inference method’s performance (Mnih & Gregor, 2014; Turner & Sahani, 2011).

For example, SVAE posterior employed here cannot directly account for the “explaining away” effect which occurs between the activations of overlapping dictionary elements (Pearl, 1988; Yu et al., 2022). Instead, “explaining away” is learned in the recognition model parameters the same way the MAP estimation in sparse coding allows for interplay between estimates under the factorial Dirac-delta posterior of sparse coding. This form of “explaining away” can indeed be seen from the simulated encoding responses in Fig. 5, capturing “extra-classical” receptive field effects such as end-stopping and contrast invariant orientation tuning (Zhu & Rozell, 2013). One important difference here is that traditional sparse coding infers all posterior parameters for small batches, allowing for more flexible explaining away if certain parameters are set to zero. Relearning the neural networks from scratch is much more computationally intensive, and thus an important next step is to build non-trivial correlations directly into the variational posteriors.

In addition to improved learning, a more complex posterior would give the recognition model the potential to exhibit interesting phenomena associated with correlations between dictionary element activations. While some computational models aim to account for such correlations (e.g., (Cadieu & Olshausen, 2009; Karklin & Lewicki, 2009; Averbeck et al., 2006)), the SVAE framework would allow for the systematic analysis of the many assumptions possible in the population coding layer within the sparse coding framework. These various assumptions can thus be validated against the population correlations observed in biological networks (e.g., (Ecker et al., 2011; Cohen & Kohn, 2011)). For example, if V1 responses are interpreted as arising via the sampling hypothesis (Fiser et al., 2010) then “explaining away” may account for correlations in neural variability observed in supra- and infra-granular layers of V1 (Hansen et al., 2012). Additionally, such correlations can be related to probabilistic population coding (e.g. (Fiser et al., 2010; Orbán et al., 2016)) where correlated variability represents correlated uncertainty in the neural code.

In this work we restricted the sparse coding model by choosing the magnitude of the output noise variance *σ*_*ϵ*_ *a priori*. This was done in order to make this work comparable to the original sparse coding implementations of (Olshausen & Field, 1996a). Nevertheless, this parameter can be fit in a data-driven way as well, providing additional performance beyond the current work. In future explorations this constraint may be relaxed.

One of the favorable properties of sparse coding using MAP inference with a Laplace prior is that truly sparse representations are produced in the sense that a finite fraction of latent variables are inferred to be exactly zero. As a consequence of the more robust variational inference we perform here, the SVAE no longer has this property. Sparse representations could be regained within the SVAE framework if truly sparse priors, which have a finite fraction of their probability mass at exactly zero, were used such as “spike and slab” priors (Garrigues & Olshausen, 2008; Ziniel & Schniter, 2013). Such sparse priors are not differentiable and thus cannot be directly incorporated into the framework presented here, however continuous approximations of sparse priors do exist and during this work we implemented an approximate spike and slab prior using a sum of Gaussians with a small and large variance respectively. We were unable in our setting, however, to replicate the superior performance of such priors seen elsewhere, and recommend additional explorations into incorporating hard-sparse priors such as the spike-and-slab or concrete (Maddison et al., 2016b) distributions.

We have restricted ourselves in this work to a relatively simple generative model of natural images. It as been noted, however, that the sparse coding model does not account entirely for the statistics of natural images (Simoncelli & Olshausen, 2001). Hierarchical variants of the sparse coding model (Karklin & Lewicki, 2003, 2005, 2009; Garrigues & Olshausen, 2010) provide superior generative models of natural images. These more complex generative models can be implemented and fit using the same methods we present here by explicitly constructing the VAE generative model in their image.

VAEs have an inherent tendency to prune unused features in the generative networks (Dai et al., 2018). Previous work has noted this effect in the case of the standard Gaussian priors. We note that similar pruning occurs in the SVAE when the priors are exponential or Cauchy as well. In these cases, the feature representation remains overcomplete, to a level of ≈1.3, which is similar to the inferred optimal overcompleteness seen in previous work (Berkes et al., 2008). Therefore the SVAE implicitly infers the overcompleteness level, a feature that previous models had to explicitly account for (Karklin & Lewicki, 2005).

The biological plausibility or our method relies on a feed-forward architecture that quickly turns input stimuli into approximate posterior distributions over the latent coefficients. While inference under this model is completely local and feed-forward, the learning through back-propagation can potentially result in more complex interactions that are at first not obviously tractable in a local neural setting. An interesting branch of work, however, aims to place backpropagation as used here and in LISTA in a biological framework (Durbin & Rumelhart, 1989; Bengio et al., 2015).

One final note is that the SVAE breaks the typical symmetry between encoder and decoder complexity. The encoder is very high-dimensional, while the decoder is a simple linear model. Despite the added complexity, we retain the ability in the SVAE to orient the encoder’s complexity towards the underlying statistics, extracting both in an unsupervised fashion. In this optimization, however, burdening the encoder with added complexity is not a concern. It is, in fact, the details of the linear decoder that matter and there are likely many local minima in the deep neural network that would help achieve similar performance under that assessment. In other statistical regimes and desired tasks, the opposite may be true (i.e., only the encoder details matter in the cost) and so flipping the complexity would be a very interesting additional path forward to expanding this philosophy further.

## 7 Conclusion

In summary, we have cast the sparse coding model in the framework of variational inference, and demonstrated how modern tools such as the VAE can be used to develop neurally plausible algorithms for inference under generative models. We feel that this work strengthens the connection between machine learning methods for unsupervised learning of natural image statistics and efforts to understand neural computation in the brain, and hope it will inspire future studies along these lines.

## Acknowledgments

The authors would like to acknowledge Ryan Pyle for involvement in early stages of this work. VG was supported by awards from the Natural Sciences and Engineering Research Council of Canada (NSERC) [PGSD3-557875-2021] and the Fonds de Recherche du Québec Nature et technologies [B2X 297667]. GB acknowledges the Marine Biological Laboratory in Woods Hole, NIMH funding for the Methods in Computational Neuroscience course (R25MH062204), and support from the Simons Foundation. JWP was supported by grants from the Simons Foundation (SCGB AWD543027), the NIH (R01EY017366), the NIH BRAIN initiative (NS104899 and 9R01DA056404-04), and the CAREER award (IIS-1150186).

## 8. Appendix

### 8.1 A Primer on Annealed Importance Sampling

To evaluate the goodness of fit we used annealed importance sampling (AIS) (R. M. Neal, 2001). Annealed importance sampling is a method to generate i.i.d. samples from an un-normalized distribution by starting with i.i.d. samples from a distribution that can be sampled from easily, and using Markov-Chain Monte-Carlo (MCMC) sampling to iteratively generate samples from distributions along a family of distributions linking the initial, known distribution to the desired distribution. It can also be used to calculate the ratio of normalizing constants for two unnormalized distributions using a family of distributions joining them. This can be used to calculate the log-likelihood by noting that

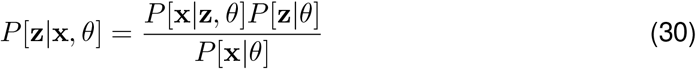

In particular, the likelihood is the factor *N* that normalizes the posterior in the expression 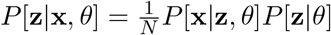.

To apply AIS, let [*p*_0_, *p*_1_, …, *p*_*N*_ ] be a family of (unnormalized) distributions such that *p*_0_ can be sampled from, *p*_*N*_ is the distribution of interest, and *p*_*n*_ and *p*_*n*+1_ are “close” to each-other in a suitable sense (for details see (R. M. Neal, 2001)). Let *M* be an MCMC method acting as our *transition operator*, such as Hamiltonian Monte Carlo (HMC, R. M. Neal (2011)) . Then, starting with a sample from *p*_0_, denoted *s*_0_, we generate a sample from *p*_1_, *s*_1_, using *s*_0_ as the initialization for MCMC, and by iterating *M* sufficiently long to generate unbiased samples from *p*_1_. We then iterate this procedure for *p*_2_ and so on until finally samples from *p*_*N*_ are generated. If *w*_*n*_ are the values of the unnormalized distributions evaluated at *s*_*n*_, then the ratio of the normalizing factors for *p*_*N*_ and *p*_0_ (denoted 𝒩_*N*_ and 𝒩_0_ respectively) can be calculated as

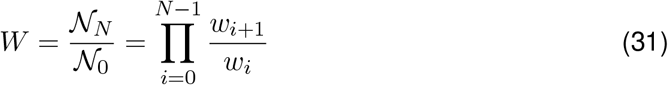

The value *W* gives an unbiased estimate of the normalizing ratio, which in the case that *p*_0_ is normalized and *p*_*N*_ is the unnormalized posterior above is equal to the likelihood. This generally leads to numerical overflow problems so we instead calculate log *W*, which in general gives a biased estimate of (a lower bound on) the log-likelihood. Averaging over many independent samples generated from AIS gives a lower-bound on the log-likelihood. The AIS method has been applied previously to evaluating goodness of fit for VAEs and other deep generative models (Wu et al., 2016).

### 8.2 Methodological Details

#### Analysis of learned basis functions

In the main text, we provide more details on the basis functions that emerge out of the SVAE inference procedure against the standard sparse coding. We analyzed the learned basis functions by fitting Gabor filters to the basis functions. Gabor filters in pixel space are Gaussian densities in Fourier space, and the orientation and frequency statistics we seek can readily be read from this density. The density peak [*f*_1_, *f*_2_] is at the spatial-(*x, y*) frequency of the filter (Gabor, 1946; Movellan, 2002), and the “frequency” and orientation metrics we report are its polar coordinate. Indeed, we report the *frequency* as the magnitude of the peak, and the *orientation* is given by the angle to the peak.

## Footnotes

1 Note that the ELBO isn’t actually increasing because, as noted above, it is always formally −∞; we could however justify this approach with a careful appeal a finite-variance *Q*(*x*) that approaches delta function only in the limit.

2 We note that this is true for digital processing only, and that analog recurrent systems can be far faster (Shapero et al., 2014).

## References

Aitchison, L., Hennequin, G., & Lengyel, M. (2018). Sampling-based probabilistic inference emerges from learning in neural circuits with a cost on reliability. arXiv preprint 1807.08952.

Averbeck, B. B., Latham, P. E., & Pouget, A. (2006). Neural correlations, population coding and computation. Nature reviews neuroscience, 7 (5), 358.

Beck, A., & Teboulle, M. (2009). A fast iterative shrinkage-thresholding algorithm for linear inverse problems. SIAM journal on imaging sciences, 2(1), 183–202.

Bengio, Y., Lee, D.-H., Bornschein, J., Mesnard, T., & Lin, Z. (2015). Towards biologically plausible deep learning. arXiv preprint 1502.04156.

Berkes, P., Turner, R., & Sahani, M. (2008). On sparsity and overcompleteness in image models. In Advances in neural information processing systems (pp. 89–96).

Berkes, P., & Wiskott, L. (2005). Slow feature analysis yields a rich repertoire of complex cell properties. Journal of Vision, 5(6), 579–602. Retrieved from http://dx.doi.org/10:1167/5.6.9

Bishop, C. M. (2005). Neural networks for pattern recognition. Oxford University Press.

Blei, D. M., Kucukelbir, A., & McAuliffe, J. D. (2016). Variational inference: A review for statisticians. arXiv preprint 1601.00670.

Boerlin, M., & Denève, S. (2011, 02). Spike-based population coding and working memory. PLoS Comput Biol, 7 (2), e1001080. Retrieved from 10.1371%2Fjournal.pcbi.1001080 doi:10.1371/journal.pcbi.1001080

Borgerding, M., Schniter, P., & Rangan, S. (2017). Amp-inspired deep networks for sparse linear inverse problems. IEEE Transactions on Signal Processing, 65(16), 4293–4308.

Boyd, S., & Vandenberghe, L. (2004). Convex optimization. Cambridge university press.

Cadieu, C., & Olshausen, B. A. (2009). Learning transformational invariants from natural movies. In Advances in neural information processing systems (pp. 209–216).

Charles, A. S., Garrigues, P., & Rozell, C. J. (2012). A common network architecture efficiently implements a variety of sparsity-based inference problems. Neural computation, 24(12), 3317–3339.

Charles, A. S., Olshausen, B. A., & Rozell, C. J. (2011). Learning sparse codes for hyper-spectral imagery. IEEE Journal of Selected Topics in Signal Processing, 5(5), 963–978.

Charles, A. S., Park, M., Weller, J. P., Horwitz, G. D., & Pillow, J. W. (2018). Dethroning the fano factor: A flexible, model-based approach to partitioning neural variability. Neural computation, 30(4), 1012–1045.

Chen, R. T., Li, X., Grosse, R. B., & Duvenaud, D. K. (2018). Isolating sources of disentanglement in variational autoencoders. Advances in neural information processing systems, 31.

Coen-Cagli, R., Dayan, P., & Schwartz, O. (2012). Cortical surround interactions and perceptual salience via natural scene statistics. PLoS Computational Biology, 8(3), e1002405.

Cohen, M. R., & Kohn, A. (2011, 6). Measuring and interpreting neuronal correlations. Nat. Neurosci., 14(7), 811–819.

Dai, B., Wang, Y., Aston, J., Hua, G., & Wipf, D. (2018). Connections with robust pca and the role of emergent sparsity in variational autoencoder models. The Journal of Machine Learning Research, 19(1), 1573–1614.

Daubechies, I., Defrise, M., & De Mol, C. (2004). An iterative thresholding algorithm for linear inverse problems with a sparsity constraint. Communications on pure and applied mathematics, 57 (11), 1413–1457.

Dayan, P., Sahani, M., & Deback, G. (2003). Adaptation and unsupervised learning. In Advances in neural information processing systems 15. MIT Press.

Doersch, C. (2016). Tutorial on variational autoencoders. arXiv preprint 1606.05908.

Durbin, R., & Rumelhart, D. E. (1989). Product units: A computationally powerful and biologically plausible extension to backpropagation networks. Neural computation, 1(1), 133–142.

Ecker, A. S., Berens, P., Tolias, A. S., & Bethge, M. (2011, 10). The effect of noise correlations in populations of diversely tuned neurons. J. Neurosci., 31(40), 14272–14283.

Fiser, J., Berkes, P., Orban, G., & Lengyel, M. (2010, March). Statistically optimal perception and learning: from behavior to neural representations. Trends Cogn. Sci. (Regul. Ed.), 14(3), 119–130.

Gabor, D. (1946, November). Theory of communication. part 1: The analysis of information. Journal of the Institution of Electrical Engineers - Part III: Radio and Communication Engineering, 93(26), 429–441. Retrieved from 10.1049/ji-3-2.1946.0074 doi:10.1049/ji-3-2.1946.0074

Garrigues, P., & Olshausen, B. A. (2008). Learning horizontal connections in a sparse coding model of natural images. In Advances in neural information processing systems (pp. 505–512).

Garrigues, P., & Olshausen, B. A. (2010). Group sparse coding with a laplacian scale mixture prior. In Advances in neural information processing systems (pp. 676–684).

Girolami, M. (2001). A variational method for learning sparse and overcomplete representations. Neural Computation, 13(11), 2517–2532.

Goris, R. L. T., Movshon, J. A., & Simoncelli, E. P. (2014, April 28). Partitioning neuronal variability. Nature Neuroscience, 17, 858. Retrieved from 10.1038/nn.3711 (Article)

Gregor, K., & LeCun, Y. (2010). Learning fast approximations of sparse coding. In Proceedings of the 27th international conference on international conference on machine learning (pp. 399–406).

Hansen, B. J., Chelaru, M. I., & Dragoi, V. (2012, November 08). Correlated variability in laminar cortical circuits. Neuron, 76(3), 590–602. Retrieved from 10.1016/j.neuron.2012.08.029 doi:10.1016/j.neuron.2012.08.029

Hubel, D. H., & Wiesel, T. N. (1962). Receptive fields, binocular interaction and functional architecture in the cat’s visual cortex. J. Physiol. (Lond.), 160, 106–154.

Jang, E., Gu, S., & Poole, B. (2016). Categorical reparameterization with gumbel-softmax. arXiv preprint 1611.01144.

Karklin, Y., & Lewicki, M. S. (2003, Aug). Learning higher-order structures in natural images. Network, 14(3), 483–499.

Karklin, Y., & Lewicki, M. S. (2005). A hierarchical bayesian model for learning nonlinear statistical regularities in nonstationary natural signals. Neural computation, 17 (2), 397–423.

Karklin, Y., & Lewicki, M. S. (2009, Jan). Emergence of complex cell properties by learning to generalize in natural scenes. Nature, 457 (7225), 83–86. Retrieved from 10.1038/nature07481

Khemakhem, I., Kingma, D., Monti, R., & Hyvarinen, A. (2020). Variational autoencoders and nonlinear ica: A unifying framework. In International conference on artificial intelligence and statistics (pp. 2207–2217).

Kingma, D. P., & Ba, J. (2014). Adam: A method for stochastic optimization. arXiv preprint 1412.6980.

Kingma, D. P., & Welling, M. (2014). Auto-encoding variational bayes. 1312.6114. Retrieved from http://arxiv.org/abs/1312.6114

Knill, D., & Richards, W. (1996). Perception as Bayesian inference. Cambridge University Press.

Knill, D. C., & Pouget, A. (2004). The bayesian brain: the role of uncertainty in neural coding and computation. TRENDS in Neurosciences, 27 (12), 712–719.

Lee, H., Battle, A., Raina, R., & Ng, A. Y. (2007). Efficient sparse coding algorithms. In Advances in neural information processing systems (pp. 801–808).

Lewicki, M., & Olshausen, B. (1999). Probabilistic framework for the adaptation and comparison of image codes. JOSA A, 16(7), 1587–1601.

Lewicki, M. S., & Olshausen, B. A. (1999). Probabilistic framework for the adaptation and comparison of image codes. JOSA A, 16(7), 1587–1601.

Maddison, C. J., Mnih, A., & Teh, Y. W. (2016a). The concrete distribution: A continuous relaxation of discrete random variables. arXiv preprint 1611.00712.

Maddison, C. J., Mnih, A., & Teh, Y. W. (2016b). The concrete distribution: A continuous relaxation of discrete random variables. CoRR, abs/1611.00712. Retrieved from http://arxiv.org/abs/1611.00712

Makhzani, A., & Frey, B. (2013). K-sparse autoencoders. arXiv preprint 1312.5663.

Martin, D., Fowlkes, C., Tal, D., & Malik, J. (2001). A database of human segmented natural images and its application to evaluating segmentation algorithms and measuring ecological statistics. In Proc. 8th int’l conf. computer vision (vol. 2, pp. 416–423).

Martins, A. F., Smith, N. A., Aguiar, P. M., & Figueiredo, M. A. (2011). Structured sparsity in structured prediction. In Proceedings of the conference on empirical methods in natural language processing (pp. 1500–1511).

Mnih, A., & Gregor, K. (2014). Neural variational inference and learning in belief networks. In Icml.

Moreno-Bote, R., Knill, D. C., & Pouget, A. (2011, Jul). Bayesian sampling in visual perception. Proc Natl Acad Sci U S A, 108(30), 12491–12496. Retrieved from 10.1073/pnas.1101430108

Movellan, J. R. (2002). Tutorial on gabor filters. Open source document, 40.

Neal, R., & Hinton, G. E. (1998). A view of the em algorithm that justifies incremental, sparse, and other variants. In Learning in graphical models (pp. 355–368). Kluwer Academic Publishers.

Neal, R. M. (2001, April). Annealed importance sampling. Statistics and Computing, 11(2), 125–139. Retrieved from 10.1023/A:1008923215028 doi:10.1023/A:1008923215028

Neal, R. M. (2011, may). Mcmc using hamiltonian dynamics. In S. Brooks, A. Gelman, G. Jones, & X.-L. Meng (eds.), Handbook of markov chain monte carlo. Chapman and Hall/CRC. Retrieved from 10.1201%2Fb10905 doi:10.1201/b10905

Olshausen, B. A. (1996). Learning linear, sparse, factorial codes.

Olshausen, B. A., & Field, D. J. (1996a, June 13). Emergence of simple-cell receptive field properties by learning a sparse code for natural images. Nature, 381, 607. Retrieved from 10.1038/381607a0

Olshausen, B. A., & Field, D. J. (1996b, May). Natural image statistics and efficient coding. Network, 7 (2), 333–339. Retrieved from 10.1088/0954-898X/7/2/014

Olshausen, B. A., & Field, D. J. (1997). Sparse coding with an overcomplete basis set: A strategy employed by v1? Vision research, 37 (23), 3311–3325.

Olshausen, B. A., & Millman, K. J. (2000). Learning sparse codes with a mixture-of-gaussians prior. In Advances in neural information processing systems (pp. 841–847).

Orbán, G., Berkes, P., Fiser, J., & Lengyel, M. (2016). Neural variability and sampling-based probabilistic representations in the visual cortex. Neuron, 92(2), 530–543.

Paszke, A., Gross, S., Chintala, S., Chanan, G., Yang, E., DeVito, Z., … Lerer, A. (2017). Automatic differentiation in pytorch.

Pearl, J. (1988). Probabilistic reasoning in intelligent systems: Networks of plausible inference. Morgan Kaufmann.

Rezende, D. J., Mohamed, S., & Wierstra, D. (2014). Stochastic backpropagation and approximate inference in deep generative models. In Proceedings of the 31st international conference on machine learning (icml-14) (pp. 1278–1286).

Ringach, D., & Shapley, R. (2004, 03). Reverse correlation in neurophysiology. Cognitive Science, 28, 147–166.

Rozell, C. J., Johnson, D. H., Baraniuk, R. G., & Olshausen, B. A. (2008). Sparse coding via thresholding and local competition in neural circuits. Neural computation, 20(10), 2526–2563.

Salimans, T. (2016). A structured variational auto-encoder for learning deep hierarchies of sparse features. arXiv preprint 1602.08734.

Sceniak, M. P., Ringach, D. L., Hawken, M. J., & Shapley, R. (1999). Contrast’s effect on spatial summation by macaque v1 neurons. Nature neuroscience, 2(8), 733.

Schwemmer, M. A., Fairhall, A. L., Denéve, S., & Shea-Brown, E. T. (2015). Constructing precisely computing networks with biophysical spiking neurons. Journal of Neuroscience, 35(28), 10112–10134.

Seeger, M. W. (2008). Bayesian inference and optimal design for the sparse linear model. Journal of Machine Learning Research, 9(Apr), 759–813.

Shapero, S., Zhu, M., Hasler, J., & Rozell, C. (2014). Optimal sparse approximation with integrate and fire neurons. International journal of neural systems, 24(05), 1440001.

Simoncelli, E. P., & Olshausen, B. A. (2001). Natural image statistics and neural representation. Annual review of neuroscience, 24(1), 1193–1216.

Smith, E. C., & Lewicki, M. S. (2006). Efficient auditory coding. Nature, 439(7079), 978.

Sun, J., Wang, X., Xiong, N., & Shao, J. (2018). Learning sparse representation with variational auto-encoder for anomaly detection. IEEE Access, 6, 33353–33361.

Theis, L., Sohl-Dickstein, J., & Bethge, M. (2012). Training sparse natural image models with a fast gibbs sampler of an extended state space. In Advances in neural information processing systems (pp. 1124–1132).

Troyer, T. W., Krukowski, A. E., Priebe, N. J., & Miller, K. D. (1998). Contrast-invariant orientation tuning in cat visual cortex: Thalamocortical input tuning and correlation-based intracortical connectivity. Journal of Neuroscience, 18(15), 5908–5927. Retrieved from http://www.jneurosci.org/content/18/15/5908 doi:10.1523/JNEUROSCI.18-15-05908.1998

Turner, R. E., & Sahani, M. (2011). Two problems with variational expectation maximisation for time-series models. In D. Barber, T. Cemgil, & S. Chiappa (eds.), Bayesian time series models (pp. 109–130). Cambridge University Press.

Tosic, I., Olshausen, B. A., & Culpepper, B. J. (2011). Learning sparse representations of depth. IEEE journal of selected topics in signal processing, 5(5), 941–952.

Van Den Oord, A., Vinyals, O., et al. (2017). Neural discrete representation learning. Advances in neural information processing systems, 30.

Velychko, D., Damm, S., Fischer, A., & Lücke, J. (2023). Learning sparse codes with entropy-based elbos.

Weiss, Y., Simoncelli, E. P., & Adelson, E. (2002). Motion illusions as optimal percepts. Nature Neuroscience, 5, 598–604.

Wu, Y., Burda, Y., Salakhutdinov, R., & Grosse, R. B. (2016). On the quantitative analysis of decoder-based generative models. CoRR, abs/1611.04273.

Yu, C., Soulat, H., Burgess, N., & Sahani, M. (2022). Structured recognition for generative models with explaining away. In S. Koyejo, S. Mohamed,

A. Agarwal, D. Belgrave, K. Cho, & A. Oh (eds.), Advances in neural information processing systems (vol. 35, pp. 40–53). Curran Associates, Inc. Retrieved from https://proceedings.neurips.cc/paper_files/paper/2022/file/003a96110b7134d678cb675c6aea6c7d-Paper-Conference.pdf

Zhu, M., & Rozell, C. J. (2013). Visual nonclassical receptive field effects emerge from sparse coding in a dynamical system. PLoS computational biology, 9(8), e1003191.

Ziniel, J., & Schniter, P. (2013). Dynamic compressive sensing of time-varying signals via approximate message passing. IEEE transactions on signal processing, 61(21), 5270–5284.

Zylberberg, J., Murphy, J. T., & DeWeese, M. R. (2011). A sparse coding model with synaptically local plasticity and spiking neurons can account for the diverse shapes of v1 simple cell receptive fields. PLoS computational biology, 7 (10), e1002250.

